# Circadian clock control of translation fidelity through MetRS-mediated methionine misincorporation

**DOI:** 10.64898/2026.06.30.735350

**Authors:** Griffin Best, Sidharth Mohan, Samuel Purvine, Deborah Bell-Pedersen

**Author notes:** Deborah Bell-Pedersen **Email:**. PNAS requires the corresponding author to provide an ORCID identifier at submission and strongly encourages all authors to use an ORCID ID. Do not include ORCIDs in the manuscript file; individual authors must link their ORCID account to their PNAS profile at www.pnascentral.org. For proper authentication, authors are not permitted to add ORCIDs on proofs. <u>Learn more or register for ORCID</u>.

## Abstract

Translation fidelity is generally viewed as a constitutive process that deteriorates under stress and aging. Here we show that the fidelity of amino acid incorporation is instead dynamically regulated by the circadian clock. In *Neurospora crassa*, methionine (Met) misincorporation into proteins exhibits robust daily rhythms, peaking at night coincident with elevated reactive oxygen species (ROS). Rhythmic Met misincorporation requires the circadian clock, the ERK-family MAPK MAK1, and MAK1-dependent phosphorylation of methionyl-tRNA synthetase (MetRS), linking circadian signaling to regulated mistranslation associated with oxidative stress resistance. Preventing MetRS phosphorylation abolishes rhythmic Met misincorporation, impairs growth, and increases sensitivity to oxidative stress, whereas a phosphomimetic MetRS mutant enhances oxidative stress survival. Proteome-wide analyses identified thousands of Met misincorporation events, including a rhythmic subset that oscillates independently of corresponding protein abundance, suggesting that mistranslation dynamically remodels proteome composition across the day. Together, these findings establish translation fidelity as a regulated circadian output and support a model in which the circadian clock temporally regulates mistranslation to enhance oxidative stress resilience.

**Significance Statement:** Biological clocks regulate translation termination fidelity, but whether they also control the accuracy of amino acid incorporation during protein synthesis was unknown. We show that the circadian clock drives rhythmic methionine misincorporation into proteins through ERK-family MAPK signaling and phosphorylation of methionyl-tRNA synthetase. Methionine misincorporation peaks during periods of elevated oxidative stress, and disrupting this regulation compromises oxidative stress survival, whereas constitutive activation enhances resistance. Together with previous work on translation termination fidelity, these findings reveal that biological clocks regulate multiple layers of translation fidelity and identify adaptive mistranslation as a mechanism that promotes cellular resilience.

## Introduction

Circadian rhythms are endogenous ∼24 h cycles that enable organisms to anticipate and adapt to predictable environmental changes, including fluctuations in light, temperature, and nutrient availability (1). In eukaryotes, these rhythms are generated by a conserved molecular oscillator composed of a transcription-translation feedback loop (TTFL) (2–4), which drives rhythmic gene expression across a large fraction of the genome (5). In *Neurospora crassa*, the core TTFL consists of the White-Collar Complex (WCC), which activates transcription of its repressor *frequency* (*frq).* FRQ protein associates with FRQ-interacting helicase (FRH) and casein kinase I (CK1) to form a repressive complex that inhibits WCC activity, completing the cycle. Progressive phosphorylation of FRQ relieves repression allowing the cycle to restart (6, 7). Through this mechanism, the TTFL generates rhythmic expression of clock-controlled genes (ccgs) that coordinate diverse physiological outputs (8, 9).

Although the TTFL operates at the level of transcription, accumulating evidence indicates that circadian regulation extends to post-transcriptional processes. Notably, about half of rhythmically accumulating proteins arise from arrhythmic mRNAs (10–13), raising the question of how temporal control of protein abundance is achieved independently of transcription. One mechanism is rhythmic mRNA translation, which has been demonstrated in *N. crassa* and other systems (11, 14–22).

In *N. crassa*, translation initiation is under circadian control through rhythmic phosphorylation of the essential initiation factor eIF2α (P-eIF2α), mediated by the CPC-3 kinase (homolog of mammalian GCN2) (15, 17). CPC-3 activity is regulated by uncharged tRNAs, whose levels oscillate, at least in part, through rhythmic aminoacylation. Consistent with this, multiple aminoacyl-tRNA synthetases (aaRSs), including ValRS, GlnRS, and AspRS are clock-controlled and peak at night (17, 23). Together, these findings support a model in which circadian regulation of aaRS activity coordinates translation initiation and protein production. In addition to regulating translation initiation, the circadian clock also controls translation termination fidelity. Rhythmic association of the ribosomal protein eL31 with translating ribosomes suppresses stop-codon readthrough during the subjective night, resulting in daily rhythms in translation termination accuracy (24). These findings suggested that the circadian clock potentially regulates multiple layers of translation fidelity, although whether it also controls amino acid incorporation fidelity was unknown.

aaRSs are essential enzymes that charge tRNAs with their cognate amino acids, ensuring accurate decoding of the genetic code during translation (25–34). Given the chemical and structural similarity among many amino acids and tRNAs, aaRSs face inherent challenges in substrate specificity and have evolved proofreading mechanisms to minimize errors (27, 28, 35, 36). Despite these safeguards, basal translation error rates can range from 0.01 - 0.1% in *Saccharomyces cerevisiae* (37) and specific mistranslation events can reach substantially higher frequencies (38).

While aaRS-mediated translation errors can be deleterious, contributing to protein misfolding, aggregation, and neurodegeneration in multiple model systems (39, 40), emerging evidence indicates that certain mistranslation events can be adaptive. In the hyperthermophilic archaeon *Aeropyrum premix*, Met misincorporation promotes adaptation of certain proteins to low temperature growth conditions (41), and in *Escherichia coli*, it enhances resistance to antibiotics (42). In *Candida albicans*, the canonical leucine CUG codon has been predominantly reassigned to specify serine, a unique deviation driven by a specialized Seryl-tRNA^CAG^ capable of pairing with the CUG codon. This tRNA is recognized by both seryl- and leucyl-tRNA synthetases, resulting in CUG being mostly translated as serine (∼97%), while a minor fraction is still read as leucine (∼3%) under standard growth conditions (43). Engineered *C. albicans* strains expressing a yeast-derived Leucyl-tRNA^CAG^ display elevated leucine incorporation at CUG sites (up to 28%), which alters cell surface protein composition and has been linked to evasion of host immune defenses during infection (43–47). This unusual codon ambiguity is thought to facilitate phenotypic variability that promotes host adaptation and pathogenicity. In mammalian cells and *S. cerevisiae*, oxidative stress activates the ERK-family of MAPKs that leads to phosphorylation of MetRS and reduces its specificity for tRNA^Met^, resulting in increased misacylation of non-cognate tRNAs and widespread Met misincorporation (48, 49). These Met residues can act as antioxidants by undergoing reversible oxidation, thereby protecting proteins and cells from ROS damage (49). These data suggested that translation fidelity is not static but may instead be regulated in response to physiological demands and environmental stress.

The circadian clock is a central regulator of cellular homeostasis and provides a mechanism for organisms to anticipate daily fluctuations in environmental and metabolic stress. In *N. crassa*, MAK1 (ERK1 homolog) activity and intracellular ROS levels exhibit circadian rhythms (50, 51). These observations led us to hypothesize that the circadian clock temporally regulates translation fidelity through MAK1-dependent phosphorylation of MetRS, driving rhythmic Met misincorporation to buffer daily oscillations in oxidative stress.

Consistent with this hypothesis, we show that Met misincorporation is clock-controlled, peaks during the night, and depends on MAK1-mediated phosphorylation of MetRS. This pathway modulates resistance to oxidative stress and extends across the proteome, where specific proteins exhibit rhythmic Met misincorporation independent of their abundance. Together, these findings reveal a mechanism by which the circadian clock dynamically regulates translation fidelity to promote cellular adaptation.

## Results

### Met misincorporation is clock-controlled in *N. crassa*

MetRS is a central determinant of Met misincorporation (48, 49, 52); therefore, we first asked whether *N. crassa* MetRS (NCU07451) expression is under circadian clock control. Analysis of published ribosome profiling (Ribo-seq) and RNA-seq data revealed that MetRS translation is rhythmic and peaked during the subjective night (17). To independently validate this, a MetRS::LUC translational reporter was generated and circadian oscillations in MetRS levels were observed that were abolished in Δ*frq* cells (**Figure S1**). These data established MetRS as a clock-controlled component of the translational machinery and positioned it as a potential contributor to rhythmic regulation of translation fidelity.

Next, to test if the circadian clock drives daily rhythms in Met misincorporation, a GFP-mCherry dual reporter system was adapted for *N. crassa* (52) (**Figure 1A**). Met at position 71 (M71) is required for mCherry fluorophore formation, and substitution of this residue abolishes fluorescence (53). Met misincorporation at position 71 in an M71 mutant restores mCherry fluorescence, and consequently, provides a readout of translation fidelity. Previous work in *S. cerevisiae* and human cell lines showed that MetRS mischarges both tRNA^Glu^ isoacceptors (tRNA^Glu(CUC)^ and tRNA^Glu(UUC)^) (48, 52), permitting Met-to-Glu substitution at M71 (mCherry^M71E^) to measure MetRS-mediated Met misincorporation. In *S. cerevisiae*, MetRS fails to misacylate tRNA^Leu(CAA)^, tRNA^Leu(GAG)^, and tRNA^Leu(UAA)^, and only weakly misacylates tRNA^Leu(UAG)^ among the four of six tRNA^Leu^ isoacceptors tested (48). Thus, an mCherry^M71L^ reporter controls for non-specific translational errors. GFP fluorescence serves as a control for translation

**Figure 1.**
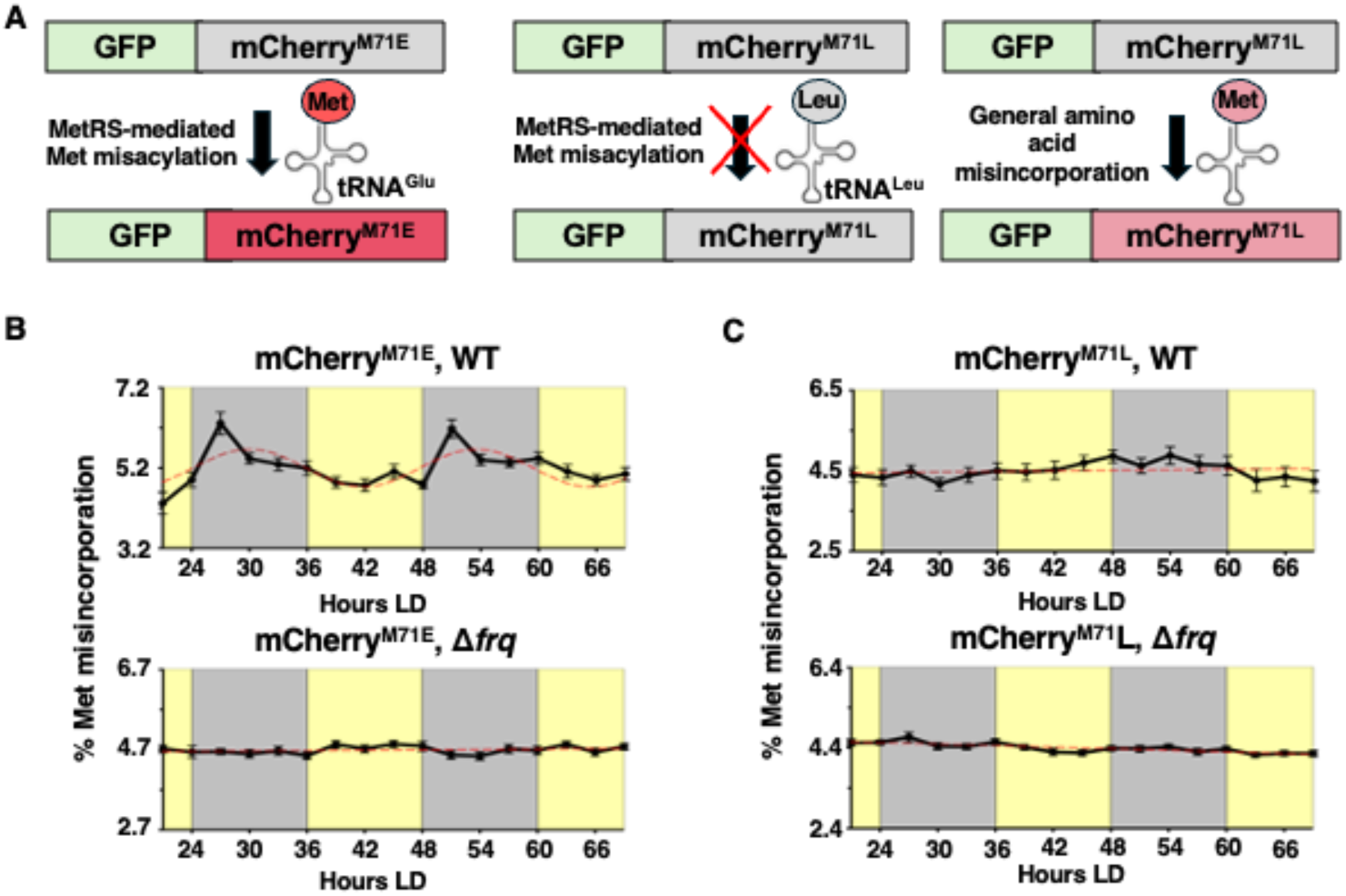
Met misincorporation in mCherry^M71E^ is clock-controlled. A) Diagram of dual-reporters. When M71 ATG is mutated to E (GAG), mCherry does not fluoresce (gray) unless tRNA^Glu^ is mischarged with Met, and Met is misincorporated (left, red). Any fluorescence from the M71 to L mutation (CTG) reflects general amino acid misincorporation of Met (right, pink) not MetRS-mediated misincorporation (middle, gray). B) Plot of the percentage of Met misincorporation into mCherry^M71E^ in WT cells grown in LD 12:12 cycles over the indicated times (Hours LD). Met misincorporation was rhythmic in the WT background as indicated by a better fit of the data to a sine wave (dotted red line; p < 0.0005) with a period of 23.5 ± 0.5 h, and arrhythmic in Δ*frq* cells as indicated by better fit to a straight line (dotted red line; p > 0.05). C) In mCherry^M71L^, Met misincorporation was low and arrhythmic in WT and Δ*frq* (dotted red line; p > 0.05). In all plots, error bars represent the mean ± SEM (n = 12). Yellow shading in B & C represents lights on and gray represents lights off.

WT and clock mutant Δ*frq* strains expressing GFP-mCherryWT (mCherryWT), GFP-mCherry^M71E^ (ATG to GAG; mCherry^M71E^), or GFP-mCherry^M71L^ (ATG to CTG; mCherry^M71L^) reporters under the copper-responsive *tcu1* promoter were generated (**Figure 1A**). The Glu codon GAG was selected based on previous studies in mammalian cells demonstrating Met misincorporation at this site (52), whereas the Leu codon CTG was selected because its frequency in the *N. crassa* genome is comparable to ATG, minimizing potential effects of codon rarity on translation efficiency and ensuring a relevant substitution context (54). In the presence of bathocuproine sulfonate (BCS), *tcu1* is constitutively active, enabling sustained reporter expression (**Figure S2A, B**). Importantly, excitation wavelengths used for fluorescence measurements did not negatively impact clock function as measured by rhythms in a FRQ::LUC translational reporter (**Figure S2C**).

In 12:12 light-dark (LD) cycles (**Figure 1**) and constant darkness (DD) (**Figure S3**), daily rhythms in Met misincorporation were observed in the mCherry^M71E^ reporter, peaking during the early night (**Figure 1B**, **S3A**), a time when MetRS levels are also elevated (**Figure S1**). These rhythms were abolished in Δ*frq* cells, indicating clock dependence. In contrast, Met misincorporation in the mCherry^M71L^ reporter was arrhythmic in both WT and Δ*frq* cells reflecting general mistakes during elongation (**Figure 1B, S3B**).

Together, these data demonstrated that the circadian clock regulates daily rhythms in amino acid incorporation fidelity under non-stress conditions and suggested that these rhythms are mediated through the MetRS misacylation pathway.

### Met misincorporation is controlled by oxidative stress, MAK1 and MetRS

Mammalian ERK MAPKs directly phosphorylate MetRS *in vitro*, and oxidative stress-induced activation of ERK signaling *in vivo* increases MetRS phosphorylation at Ser209 and Ser825 promoting Met misincorporation (49, 52). Because the ERK1 homolog MAK1 is clock-regulated in *N. crassa* (50), we tested whether MAK1 activity contributes to MetRS phosphorylation and Met misincorporation.

To directly assess this pathway, we examined phosphorylation of MAK1 under oxidative stress. P-MAK1 levels increased significantly following hydrogen peroxide (H_2_O_2_) treatment, whereas the ERK-family MAPK P-MAK2 levels did not (**Figure S4**), supporting that MAK1 is the primary kinase responding to oxidative stress in this context.

Oxidative stress, which activates ERK signaling, was induced using H_2_O_2_ (49). In MetRS WT mCherry^M71E^ reporter strains, Met misincorporation increased significantly at 100 mM H_2_O_2_ (**Figure 2A, S4C**), whereas no effect was observed in the MetRS WT mCherry^M71L^ control (**Figure 2B, S4D**). These results were consistent with prior studies showing ERK-dependent increases in Met misincorporation (52) and support a role for MAK1-MetRS signaling in regulating translation fidelity.

**Figure 2.**
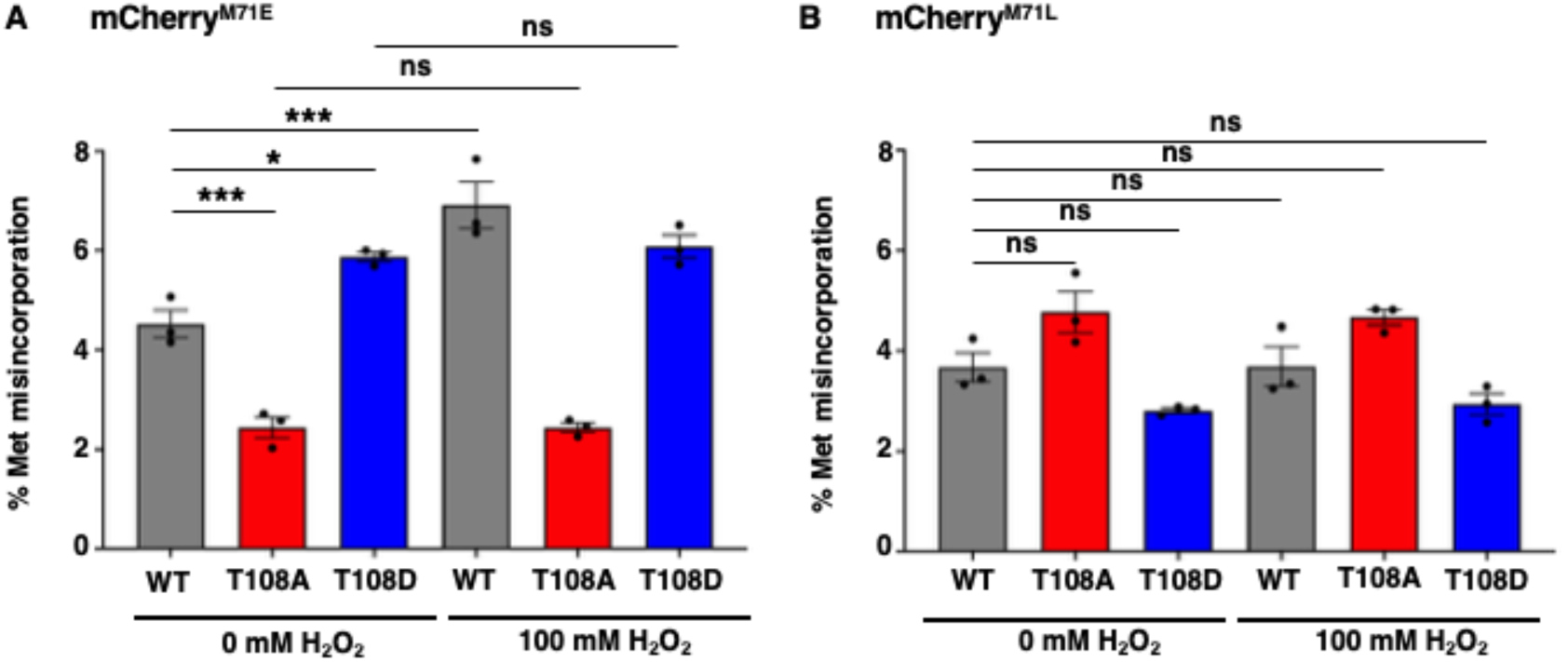
Oxidative stress increases Met misincorporation through phosphorylation of MetRS. Plots of percent Met misincorporation in WT, *metRS*^T108A^ (T108A), and *metRS*^T108D^ (T108D) cells expressing A) mCherry^M71E^ or B) mCherry^M71L^ and grown in DD at 25 °C with 0 mM or 100 mM H_2_O_2_ to induce oxidative stress. Measurements of Met misincorporation were taken five days post-H_2_O_2_ treatment (DD 114). Error bars represent the mean ± SEM (n = 3; *p < 0.05, **p < 0.005, ***p < 0.0005), and ns represents no significant change.

To identify potential ERK-family MAPK-dependent phosphorylation sites on *N. crassa* MetRS, a MetRS polyclonal antibody was generated for immunoprecipitation (**Figure S5**) followed by phosphoproteomics. A single phosphorylation site, Threonine108 (T108), was detected specifically in WT cells, but not Δ*mak1* cells (**Dataset S1**). T108 is immediately followed by a proline (T108-P109), consistent with the minimal ERK-family MAPK recognition motif (55). Importantly, T108 phosphorylation was detected in Δ*mak2* cells, supporting MAK1 as the kinase responsible for directing MetRS phosphorylation.

To test the functional significance of this site, phospho-null (*metRS*^T108A^) and phosphomimetic (*metRS*^T108D^) mutants were generated. MetRS abundance was unchanged in both mutants (**Figure S6**). In the mCherry^M71E^ reporter strains, the overall levels of Met misincorporation were reduced in *metRS*^T108A^ and increased in *metRS*^T108D^ compared to the WT control (0 mM H_2_O_2_), and both mutants were insensitive to oxidative stress (100 mM H_2_O_2_) (**Figure 2A**). Notably, these mutations had little effect on the mCherry^M71L^ reporter with or without oxidative stress (**Figure 2B**). These results demonstrated that MAK1-dependent phosphorylation of MetRS at T108 modulates Met misincorporation in response to oxidative stress.

### MetRS phosphorylation drives rhythmic Met misincorporation

To determine if MAK1-dependent phosphorylation of MetRS underlies circadian rhythms in Met misincorporation, Met misincorporation was assayed in Δ*mak1* and the *metRS*^T108^ mutants in L 30 °C/D 25 °C cycles. These conditions were used to assay rhythmicity in the mutants because light and temperature entrainment reinforced circadian amplitude and synchrony across the population, enabling more robust detection of Met misincorporation rhythms than in DD 25 °C.

Using the mCherry^M71E^ reporter, Met misincorporation was low and arrhythmic in the *metRS*^T108A^ mutant, and high and arrhythmic in the *metRS*^T108D^ mutant (**Figure 3A**). Both mutants were arrhythmic in mCherry^M71L^ with Met misincorporation levels similar to WT (**Figure 3B**). However, the Δ*mak1* strain showed low and arrhythmic misincorporation in both mCherry^M71E^ and mCherry^M71L^ reporters (**Figure 3A, B**), suggesting that MAK1 regulates both MetRS-dependent and MetRS-independent mechanisms influencing Met misincorporation. Together, these findings demonstrated that MAK1-dependent phosphorylation of MetRS at T108 is required for circadian control of Met misincorporation.

**Figure 3.**
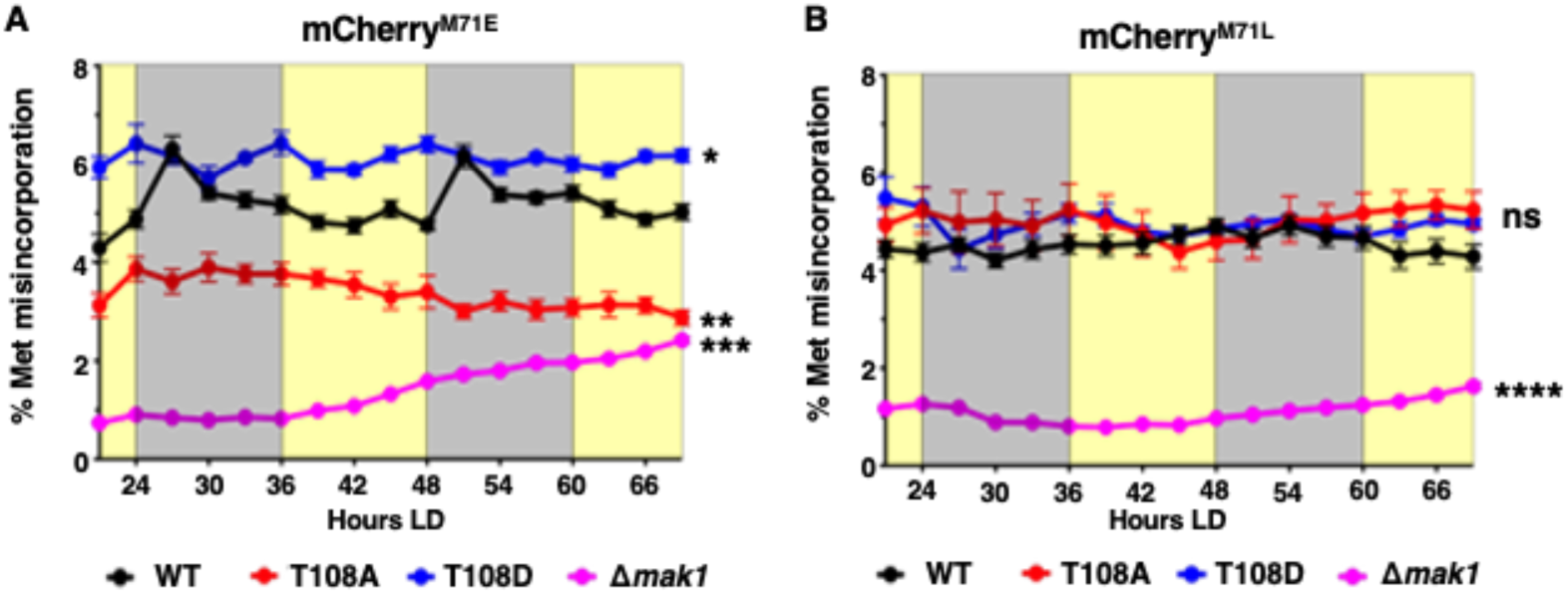
MAK1-dependent phosphorylation of MetRS is essential for Met misincorporation rhythms. Plot of the percentage of Met misincorporation into A) mCherry^M71E^ or B) mCherry^M71L^ B) in WT (black), *metRS*^T108A^ (T108A, red), *metRS*^T108D^ (T108D, blue) and Δ*mak1* (pink) cells. The experiments were conducted with those shown in Figure 1 B & C, and the WT data is re-plotted here. The plots are as described in Figure 1. mCherry^M71E^ Met misincorporation levels in *metRS*^T108A^ and Δ*mak1* cells were significantly lower, whereas Met misincorporation levels in *metRS*^T108D^ were significantly higher than WT, and Met misincorporation rhythms were abolished in all 3 mutants. mCherry^M71L^ Met misincorporation levels in *metRS*^T108A^ or *metRS*^T108D^ cells were not significantly different from WT; however, Met misincorporation in Δ*mak1* cells was significantly decreased. Error bars represent the mean ± SEM (n = 12; **p < 0.005, ***p < 0.0005, ****p < 0.0001), and ns represents no significant change.

### MetRS-mediated Met misincorporation promotes resistance to oxidative stress

Met misincorporation was shown to protect HeLa cells from oxidative damage (49). Because ROS levels peak during the night in *N. crassa* (51), coinciding with maximal Met misincorporation, we hypothesized that this process enhances stress resistance.

Cell viability assays revealed that exposure to 100 mM H_2_O_2_, but not 0 mM H_2_O_2,_ caused progressive cell death in WT, Δ*frq*, and *metRS*^T108D^ cells grown in DD, while the *metRS*^T108A^ mutant was significantly more sensitive and lost viability rapidly (**Figure 4A,B**). At intermediate stress (50 mM H_2_O_2_), *metRS*^T108D^ mutant cells displayed enhanced survival, and *metRS*^T108A^ mutants sustained reduced viability, relative to WT and Δ*frq* cells (**Figure 4C**), consistent with increased Met misincorporation conferring protection against oxidative stress. No viability defects were observed in WT cells grown in 10 mM H_2_O_2_ (**Figure S7**), correlating with minimal increases in Mak1 phosphorylation and Met misincorporation at this dose (**Figure S4**). Importantly, blocking MetRS phosphorylation also significantly reduced growth rate even in the absence of applied oxidative stress, with the *metRS*^T108A^ mutant showing the strongest impact (**Figure 4D**). To determine whether this phenotype reflected a general defect in gene expression or protein synthesis, GFP accumulation from the dual reporter system was quantified. GFP levels were similar in WT, *metRS*^T108D^, and *metRS*^T108A^ mutant strains over the time course (**Figure 4E**), indicating that disruption of MetRS phosphorylation does not cause a major defect in reporter expression or overall translational capacity. These findings suggest that the reduced growth of *metRS*^T108A^ and *metRS*^T108D^ cells is more likely due to loss of regulated Met misincorporation rather than a global impairment of protein production. Together, these findings support a model in which MAK1-dependent phosphorylation of MetRS promotes adaptive Met misincorporation that enhances resistance to oxidative stress in a dose-dependent manner. Furthermore, the reduced growth of *metRS*^T108A^ and *metRS*^T108D^ mutant cells under non-stress conditions suggested that circadian regulation of the MetRS mistranslation pathway contributes to cellular fitness even in the absence of externally applied oxidative stress, potentially by anticipating daily fluctuations in endogenous metabolic or oxidative stress.

**Figure 4.**
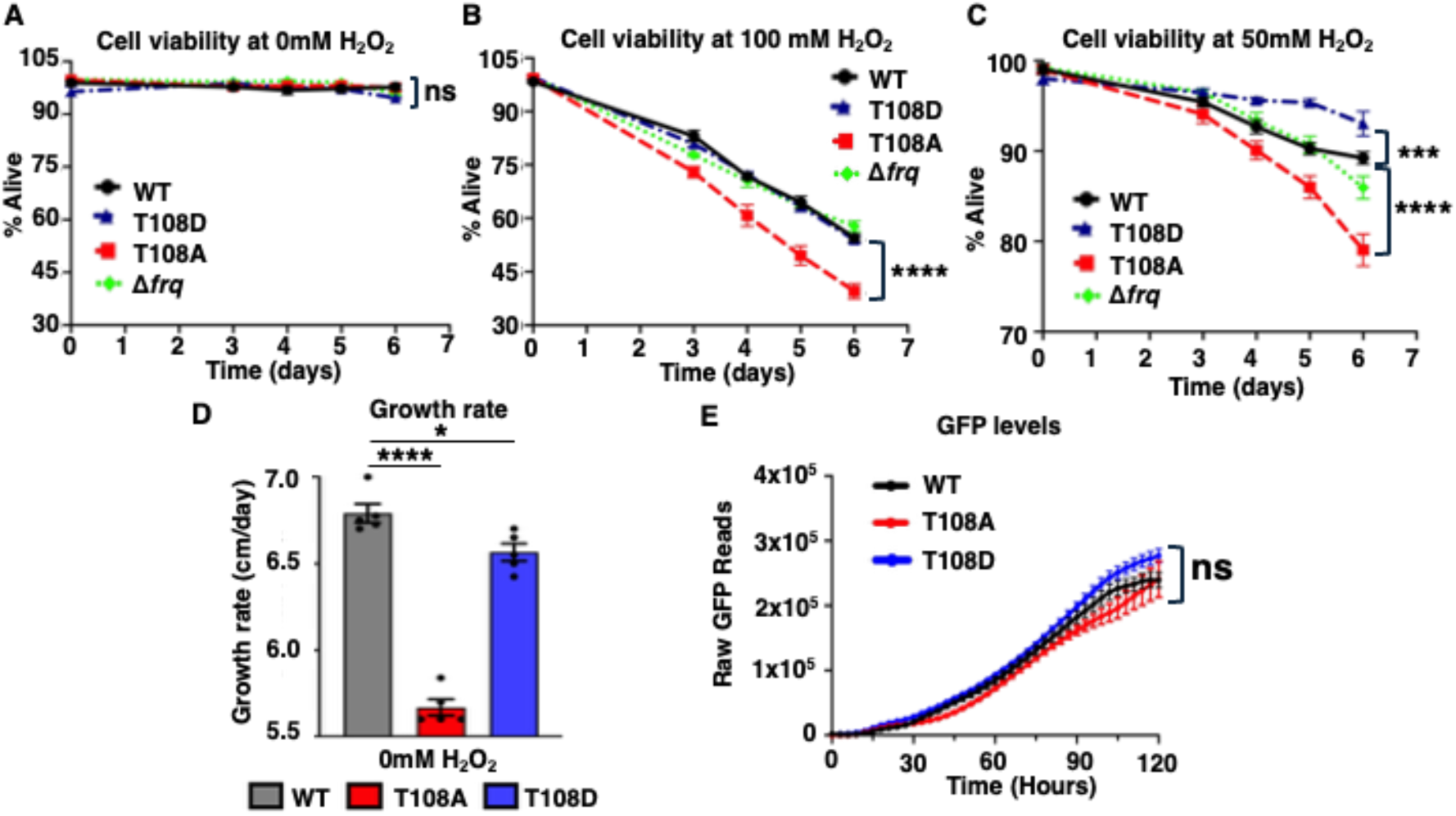
Cell viability is altered in *metRS*^T108A^ and *metRS*^T108D^ mutants following oxidative stress. Plots of cell viability for WT (black), *metRS*^T108A^ (T108A, red), *metRS*^T108D^ (T108D, blue) and Δ*frq* (green) cells grown in A) 0 mM, B) 100 mM, and C) 50 mM H_2_O_2_. Cell viability was measured at zero-, three-, four-, five-, and six-days post-treatment. *metRS*^T108A^ cells were less viable than all groups when grown in 50 or 100 mM H_2_O_2_, and *metRS*^T108D^ cells were more viable than all groups when grown in 50 mM H_2_O_2_. D) Preventing MetRS phosphorylation impairs growth. Plot of growth rate of WT (black), *metRS*^T108A^ (T108A, red), and *metRS*^T108D^ (T108D, blue) mutant cells grown in DD. E) GFP accumulation from the dual reporter system in WT (black), *metRS*^T108A^ (T108A, red), and *metRS*^T108D^ (T108D, blue) strains. GFP levels of the mutants were not significantly different from WT, indicating that altered MetRS phosphorylation does not substantially affect reporter expression or overall protein production. Error bars represent the mean ± SEM (n = 4; ***p < 0.0005, ****p < 0.0001, ns represents no significant change).

### The *N. crassa* proteome exhibits rhythmic Met misincorporation

To determine if clock control of Met misincorporation occurs *in vivo* under constant growth conditions, a published circadian proteomics dataset from *N. crassa* was analyzed using Andromeda (10, 56). Broadly, 2,637 unique Glu to Met and 1,278 Asp to Met substitutions were identified across all time points, representing a marked enrichment compared to most other Met misincorporation events (**Figure 5; Dataset S2 and S3**). For example, Leu to Met substitutions were far less frequent (299 events) despite the higher genomic abundance of Leu codons. These findings are consistent with selective enrichment for Met misincorporation at Glu and Asp codons (48). Analysis of the codons underlying Met substitution events revealed non-uniform representation among synonymous codons, particularly for Glu to Met substitutions (**Table S1**). For Glu-to-Met substitutions, GAG was more frequently detected than GAA, whereas Asp-to-Met substitutions were more common at GAC than GAT, consistent with previous studies in yeast (48). Leu-to-Met substitutions were distributed across all six Leu codons, with CTC the most frequently detected. These findings suggest that codon identity, tRNA abundance, or local sequence/protein context may influence the accumulation or detection of Met misincorporation events.

**Figure 5.**
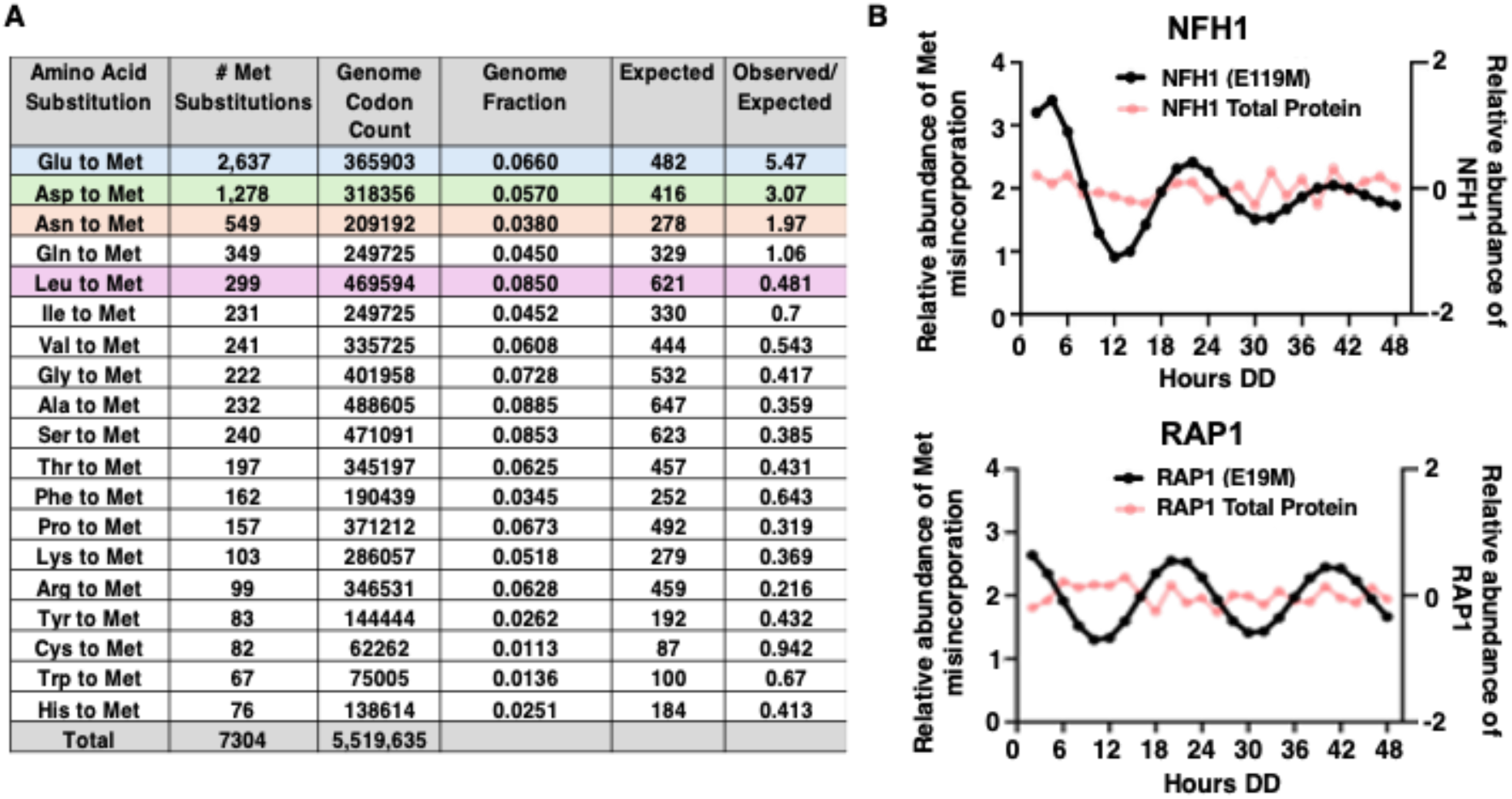
Specific proteins undergo rhythmic Met misincorporation *in vivo*. A) Glu to Met (blue), as well as Asp (green) and Asn (orange) misincorporation is enriched compared to Leu to Met (pink). Genome fraction (column 4) is calculated as the codon count (column 3) divided by the sum of all codons (5,519,635). The expected number of Met substitutions (column 5) is calculated by multiplying the genome fraction (column 4) by the number of observed Met substitutions (column 2) B) ECHO-fitted data of Met misincorporation at the indicated amino acids for two representative proteins (black) over time (Hours DD) (left axis). Total protein levels (pink) from the same dataset (unfitted data) are shown (right axis).

To obtain more robust Glu to Met peptide quantifications, the Met misincorporated peptide search was redone using MS-GF+ (10). In this analysis, there was enrichment of Glu to Met (1554 events) compared to Leu to Met (203 events) substitutions, validating the Andromeda search data (**Dataset S4**). The Glu to Met peptides were tested for rhythmicity using the R-program ECHO (57), which identified 236 peptides from WT *N. crassa* cells with oscillating Met misincorporation (**Dataset S5**). After excluding proteins with rhythmic abundance from the same published data set (10), 130 unique peptides remained that exhibited rhythms in misincorporation independent of rhythms in the corresponding protein with two examples shown (**Figure 5B)**. In these examples, Met misincorporation peaked during the subjective night coinciding with the mCherry^M71E^ reporter in WT strains (**Figure 1B**). Overall, a majority (59%) of rhythmic Met misincorporation events peaked during the subjective night (**Dataset S5**). A low confidence Met misincorporation event was identified in the core clock protein WC-1, but not WC-2, FRQ, or FRH (**Dataset S3**). Consistent with limited effects on the core oscillator, the *metRS*^T108A^ mutant displayed only a modest alteration in the period of FRQ::LUC rhythms and no detectable defects in developmental rhythms compared to WT (**Figure S8**). Gene ontology analysis of proteins exhibiting Met misincorporation revealed enrichment for pathways associated with oxidative stress responses, translation, and central carbon metabolism (**Dataset S5**), consistent with a role for rhythmic mistranslation in coordinating metabolic and stress-responsive processes. Together, these results demonstrated that the circadian clock drives protein-specific rhythms in Met misincorporation across the proteome.

## Discussion

The *N. crassa* circadian clock regulates translation termination fidelity through stop-codon readthrough (SCR) (24). Ribosomal protein eL31 associates with ribosomes rhythmically, peaking during the subjective night where it suppresses SCR and enhances termination fidelity. Consequently, SCR is elevated during the day when eL31 occupancy is reduced. In parallel, our findings show that the clock also modulates amino acid incorporation fidelity through Met misincorporation. Together, these studies support that the circadian clock orchestrates multiple layers of translation fidelity, dynamically modulating translation accuracy in response to time-of-day-dependent cellular demands and stress.

Our data identify a mechanism by which the circadian clock regulates amino acid incorporation fidelity through rhythmic Met misincorporation. MAK1-dependent phosphorylation of MetRS at T108 drives daily rhythms in Met misincorporation that peak during the night (**Figure 1**), coinciding with elevated MetRS abundance (**Figure S1**). Together, these findings suggest that circadian regulation of MetRS expression and phosphorylation are temporally coordinated to modulate translation fidelity. Although we did not directly test whether MAK1 phosphorylates MetRS *in vitro*, several observations support this model. ERK-family MAPKs directly phosphorylate MetRS in mammalian cells, where phosphorylation induces a conformational change that broadens tRNA recognition and promotes misacylation of non-cognate tRNAs (49). Consistent with a conserved mechanism, MetRS T108 phosphorylation was detected in WT and Δ*mak2* cells following oxidative stress but was absent in Δ*mak1* cells, and disruption of either MAK1 or the MetRS T108 phosphosite abolished rhythmic Met misincorporation. Future biochemical studies will be needed to determine whether MetRS T108 phosphorylation similarly alters the substrate specificity of *N. crassa* MetRS.

Oxidative stress activates MAK1-dependent phosphorylation of MetRS, reducing fidelity and increasing Met misincorporation (**Figure 2**), and disruption of this pathway through T108 mutations or *mak1* deletion abolishes rhythmic Met misincorporation (**Figure 3**). Functionally, modulation of this pathway alters oxidative stress sensitivity (**Figure 4**): phosphomimetic *metRS*^T108D^ cells exhibited enhanced survival under intermediate oxidative stress, whereas phospho-null *metRS*^T108A^ cells showed reduced viability under both intermediate and high oxidative stress conditions. Importantly, *metRS*^T108A^ and *metRS*^T108D^ cells also exhibited reduced growth even in the absence of applied oxidative stress (**Figure 4**). Because MetRS mutant cells retained normal viability, normal MetRS abundance, and accumulated GFP from the dual reporter at levels similar to WT (**Figure 4E**), this phenotype is unlikely to reflect a general defect in transcription, translation initiation, or overall protein synthesis. Instead, the selective effects of MetRS-dependent mistranslation reporters indicate that loss of regulated Met misincorporation contributes to the growth defect. These findings suggest that circadian regulation of MetRS-dependent mistranslation contributes to cellular fitness under basal conditions, potentially through the generation of protein variants that help cells anticipate endogenous metabolic and oxidative stress. Although broader effects of the MetRS mutants on MetRS activity cannot yet be excluded, the selective effects on MetRS-dependent mistranslation reporters together with unchanged MetRS abundance indicate a role for the MetRS mutants regulating substrate specificity. The requirement for MAK1 and the T108 phosphosite for rhythmic misincorporation strongly supports temporal regulation of this modification. Together, these findings suggest a model in which the circadian clock temporally tunes amino acid incorporation fidelity to anticipate daily fluctuations in oxidative stress.

While our data identify MetRS T108 phosphorylation as a key regulator of rhythmic Met misincorporation, deletion of *mak1* reduced Met misincorporation more strongly than the *metRS*^T108A^ mutation and affected both MetRS-dependent (mCherry^M71E^) and independent (mCherry^M71L^) reporters (**Figure 3**). These findings suggest that MAK1 regulates Met misincorporation through additional mechanisms beyond MetRS phosphorylation. One possibility is regulation of elongation dynamics. In mammalian systems, ERK signaling modulates eEF2 kinase (eEF2K), and in *Caenorhabditis elegans*, eEF2 phosphorylation slows translation elongation and enhances amino acid selection fidelity under some conditions (58, 59). Consistent with this we previously showed that eEF2 phosphorylation is clock-controlled and peaks during the day, coinciding with reduced translation rates and increased fidelity (14). Together, these observations support the idea that MAK1 coordinates multiple layers of translation fidelity control, including both amino acid selection and elongation dynamics. However, if MAK1 acts through multiple translation fidelity pathways, an important question is why rhythms are not observed in the mCherry^M71L^ reporter? A likely explanation is that the GFP-mCherry dual reporter system lacks the sensitivity to detect low-amplitude rhythms independent of MetRS activity. This limitation highlights the need for more sensitive and robust approaches to fully resolve circadian regulation of amino acid incorporation fidelity across the proteome.

An important nuance emerging from these data is the relationship between ROS, MAK1 activity, and circadian timing. While ROS can activate MAK1 (**Figure S4**), it is unlikely to be the primary driver of MAK1 rhythmicity. ROS levels peak during the night, whereas P-MAK1 levels peak during the afternoon (50, 51), suggesting that circadian regulation of MAK1 is driven by upstream clock-controlled pathways, with ROS acting as a modulatory input. These observations suggest that basal circadian regulation establishes the temporal profile of MAK1 activity, while ROS provides an acute modulatory signal that further enhances pathway activation under stress. Because Met residues can function as ROS scavengers, rhythmic Met misincorporation could itself contribute to daily fluctuations in cellular ROS levels by increasing the abundance of oxidation-sensitive Met residues at specific times of day. In this model, circadian regulation of MetRS activity would not only respond to oxidative stress but could also impact temporal control of cellular redox homeostasis.

The magnitude of increased Met misincorporation at night compared to the day is modest (∼2%) (**Figure 1B**), but this likely underestimates the overall impact of this pathway. Our reporter specifically captures Glu to Met substitutions at GAG codons, whereas previous studies indicated that MetRS can also mischarge tRNA^Asp^ and tRNA^Lys^ (48). Thus, rhythmic Met misincorporation may extend to multiple amino acid contexts, potentially broadening its proteome-wide effects. Consistent with this idea, mass spectrometry analysis also revealed enrichment of Asp to Met substitutions, although not Lys to Met, in *N. crassa* (**Figure 5A**). Expanding the reporter system to additional residues will be important for capturing the full scope of circadian regulation of amino acid incorporation fidelity.

The frequency of Met misincorporation detected by the GFP-mCherry reporter is not directly comparable to the number of Met-substituted peptides identified by mass spectrometry. The reporter provides a targeted readout of a permissive Glu to Met substitution under controlled expression conditions, whereas proteomic detection of mistranslation events is limited by peptide abundance, ionization efficiency, stochastic peptide sampling, and stringent filtering thresholds (60). In addition, some mistranslated proteins may be rapidly degraded or fall below detection limits. Thus, the proteomics dataset likely represents only a subset of total Met misincorporation events occurring across the proteome. The mechanisms that determine which Met misincorporation events accumulate in the proteome remain unclear. One possibility is that substitutions occurring at solvent-exposed residues are selectively retained because they enhance ROS buffering, as proposed previously (49). Alternatively, Met misincorporation may occur more broadly, with substitutions at structurally sensitive positions resulting in protein misfolding and degradation. Under this model, the proteome would represent a filtered subset of tolerated Met misincorporation events. Similarly, only a subset of Met-misincorporated peptides exhibited detectable circadian rhythms, likely reflecting the combined constraints of proteomic sampling depth, protein turnover, and the requirement for sufficient rhythmic amplitude to achieve statistically-supported rhythmicity. Distinguishing between selective retention of beneficial substitutions and proteostasis-mediated removal of deleterious variants will be an important goal of future studies.

Proteomic analysis demonstrated that Met misincorporation occurs broadly across the proteome, with a subset of proteins exhibiting circadian rhythms independent of protein abundance (**Figure 5 and Dataset S2-S5**). These data raise the possibility that rhythmic Met misincorporation contributes to temporal regulation of protein function. Consistent with this idea, proteins exhibiting rhythmic Met misincorporation were enriched for pathways linked to oxidative stress responses and metabolism. Although we have not yet directly tested the functional consequences of misincorporation events at specific sites, even low-frequency events could have significant effects if they occur at regulatory or structurally sensitive residues. More generally, because Met residues can influence protein structure, stability, and localization, and can act as reversible redox buffers, rhythmic substitution at defined sites may tune protein activity in a time-of-day-dependent manner.

Collectively, our findings support that translation fidelity itself is a regulated circadian output. In addition to Met misincorporation and SCR, other fidelity mechanisms may be clock-controlled. For example, oxidative stress can induce mistranslation through other aaRSs, such as ThrRS-mediated tRNA^Ser^ mischarging (61). Furthermore, the relative abundance of aaRSs and their cognate tRNAs can influence fidelity (62), and many aaRSs exhibit circadian expression (15, 23). These findings add to growing evidence that aaRSs function not only as core translational machinery, but also as dynamically regulated signaling-responsive determinants of proteome composition and translation fidelity.

Finally, these findings have important implications for aging. Both circadian rhythms and translation fidelity decline with age and are linked to proteostasis defects and age-related disease (63–67). Our results suggest a mechanistic connection between these processes, raising the possibility that age-associated damping of circadian amplitude leads to reduced regulation of translation fidelity. Notably, ERK signaling also declines with age (68, 69), positioning it as a potential integrator of circadian and proteostasis pathways. Future studies should investigate whether restoring circadian amplitude can preserve translation fidelity and improve cellular resilience during aging.

In summary, this work reveals a mechanism by which the circadian clock dynamically regulates translation fidelity through MetRS-dependent mechanisms. By temporally modulating amino acid incorporation fidelity, the clock may tune proteome composition and stress resilience to anticipate predictable daily fluctuations in oxidative state.

## Materials and Methods

### *N. crassa* strains and growth conditions

*N. crassa* vegetative growth conditions, transformation, crossing protocols, and race tube protocols were as described previously (70, 71). Details of strain construction are provided in **SI Methods**, and strains, key reagents, and oligonucleotide primers used in this study are listed in **Table S2**.

### Oxidative stress to induce ERK-family MAPK phosphorylation and for test of dual reporters

To induce oxidative stress for MAK1/2 phosphorylation, tissue (mycelial mats) was treated with H_2_O_2_ at a final concentration of 0, 10 and 100 mM for 5 min before harvesting. Flash-freezing of tissue in liquid N_2_ and grinding was as previously described (19).

To induce oxidative stress of the dual reporter strains (mCherryWT, mCherry^M71E^, and mCherry^M71L^), germinated conidia were grown in constant light (LL) at 30 °C for six days in 1X Vogels minimal media with 2% glucose (1X V2G) and 5 μM CuSO_4_ to inhibit basal expression of the dual reporter. Conidia were collected using 1X V2G and filtered through cheesecloth to remove excess mycelium. Next, conidia were pelleted at 3000 rpm for three minutes and suspended in 500 μL of 1X V2G. Cell concentration was measured using a spectrophotometer (λ=420) at a 1:100 dilution, and cells were diluted to 1 X 10^7^ cells/mL (assuming OD_420_ of 1 = 5 X 10^6^ cells/mL). Revvity Optiplate-96 plates were prepared with 150 μL of 1X V2G, BCS, and H_2_O_2_. 100 μL of cells (1,000,000 cells) were plated resulting in a final volume of 250 μL and final concentrations of 200 μM BCS and 0 mM, 10 mM, or 100 mM of H_2_O_2_. Plates were covered with a Breathe-Easy membrane and moved to an EnVision plate reader with a monochromator in DD at 25 °C. mCherry and GFP excitation wavelengths (578 nM and 484 nM, respectively) were applied, and emission wavelengths (618 nM and 517 nM, respectively) were measured every three hours over five days. Percent Met misincorporation was calculated by taking the ratio of mCherry (mCherry^M71E^ or mCherry^M71L^) to GFP signal over the ratio of mCherryWT to GFP signal and then multiplied by 100 to give a final percent.

### Time courses with dual reporters and FRQ::LUC

Dual-reporter (GFP and mCherry) strains were grown in LL at 30 °C for six days and purified as described above (Oxidative stress test with dual reporters), with one exception: cells were grown in 1X V2G without 5 μM CuSO_4_. Revvity Optiplate-96 plates with agar were prepared with 200 μM BCS as previously described (15) with one exception, 200 μM BCS replaced 10 μΜ luciferin. 500 cells were seeded onto plates and incubated in LL at 30 °C for one day before moving them to an EnVision plate reader with a monochromator in L (30 °C)/D (25 °C) cycles or DD at 25 °C. Measurements of FRQ::LUC rhythms were performed as previously described (15).

### Generation of anti-MetRS antibody

To detect MetRS protein in western blots, MetRS was tagged with V5. However, the *metRS::V5* strain (DBP3953) had a growth defect, suggesting altered MetRS function. Therefore, we generated an antibody against *N. crassa* MetRS. The full *N. crassa* MetRS protein was fused to a cysteine protease domain (CPD) and ligated into an *E. coli* expression vector (pET22b(+)). Bacterial protein expression and purification were as previously described (72) with one exception, protein-bound Ni-NTA beads were incubated with 600 μL of lysis buffer containing Phytic acid (InsP_6_) (833 μg/mL) and shaken on ice for 1.5 hours to cleave the CPD domain from MetRS (NCU07451). After purification by fast protein liquid chromatography (FPLC), purified MetRS was submitted to the Pocono Rabbit Farm & Lab (Canadensis, PA) where rabbits were immunized and subsequently bled for anti-MetRS antibody serum.

### Protein extraction and western blots of MetRS and P-MAK1/2

Protein extraction and western blot protocols were as previously described (16). Anti-MetRS antibody was diluted at a 1:10,000 in 5% non-fat milk in 15mL 1X TBST. Anti-P-MAK1/2 (anti-P-p44/42) and anti-MAK2 (anti-p44/42) antibodies were diluted at a 1:1,000 in 5% non-fat milk in 15mL 1X TBST. Anti-V5 antibodies were diluted 1:5000 in 5% non-fat milk in 15mL 1X TBST. Western blots were analyzed using an Odyssey-CLX infrared imaging system (Li-Cor).

### Identification of peptides with Met substitutions

Mass spectrometry of proteins isolated from pooled WT and Δ*csp1* cells over a circadian time course (10) were reanalyzed for Met substitutions in peptides using MaxQuant (v2.6.7.0) (73, 74) with the integrated Andromeda search engine (56). Amino acid substitutions with Met were included as variable modifications in the database search. Peptides shorter than 7 amino acids were excluded from the search to obtain reliable peptide-to-protein mapping. To reduce computational complexity and false positives, a maximum of 3 misincorporation events per peptide were allowed during the search for each amino acid. Site assignments correspond to the highest-scoring localization reported by MaxQuant, and corresponding localization probabilities for all alternative sites are provided in **Dataset S2**. MS-GF+ analysis on characterized WT pools was performed using a previously described workflow (10). To identify Glu to Met and Leu to Met peptides, the search done using annotation lists where every Glu or Leu in the *N. crassa* coding sequence was replaced with Met. Care was taken to ensure substituted-peptide identifications were from previously non-filter passing tandem mass spectra and do not replace any identifications from the previous data. All Met misincorporated peptides identified by MS-GF+ analyses are provided in **Dataset S4**. ECHO was used to identify rhythmic Glu to Met substitutions in WT samples as described (57).

### Cell viability assays

Strains and Revvity Optiplates-96 plates were prepared as described for oxidative stress experiments. Following seeding, the plates were incubated in DD at 25 °C, and viability was measured on days three, four, five, and six. Cells were resuspended and mixed 1:1 (8 μL) with 0.4% Trypan blue and then incubated in DD for 5 min. Subsequently, 8 μL of the mixture was loaded onto a hemocytometer, and both live (clear) and dead (blue) cells were counted by light microscopy to determine the percentage of viable cells.

### Statistical analysis

Circadian time course data were analyzed using F-tests to compare the fit of the data to a sine wave or a line, as previously described (50, 70). Student’s t-tests were used to assess differences in P-MAK1 and total MetRS levels. P-MAK1 levels were normalized to total MAK-2 levels, and MetRS levels were normalized to total protein levels (Amido Black staining). Student’s t-tests were also used to determine differences in *ras-1^bd^*period. Simple linear regression was applied to cell viability data. Repeated measurements two-way ANOVA was used to assess oxidative stress data and time course comparisons among WT, Δ*frq*, *metRS*^T108A^, and *metRS*^T108D^ strains. All statistical analyses were performed using GraphPad Prism with at least three biological replicates.

## Supporting information

Supporting text: Detailed Strain Construction Figures S1 to S7 Tables S1 to S2 Legends for Datasets S1 to S5 SI References

## Acknowledgments

We thank Drs. Teresa Lamb and Christopher Weitzel for insightful input into this work, and the Fungal Genetics Stock Center for strains used this study. Funding for this study was provided by NIH R35 GM126966 grant to DBP.

## Author Contributions

Conceptualization, D.B.P. and G.B., Investigation, G.B., S.M. and S.P., Data Analyses and Visualization, G.B., Writing, G.B. and D.B.P., Funding Acquisition, D.B.P.

## Competing Interest Statement

The authors declare no competing interests.

## Notes

https://figshare.com/s/89a474f54c67f67be215

### Competing Interest Statement

The authors have declared no competing interest.

