## Supporting text: Detailed Strain Construction Figures S1 to S7 Tables S1 to S2 Legends for Datasets S1 to S5 SI References for "Circadian clock control of translation fidelity through MetRS-mediated methionine misincorporation"

PNAS

**This PDF file includes:**

Supporting text: Detailed Strain Construction
Figures S1 to S7
Tables S1 to S2
Legends for Datasets S1 to S5
SI References

**Other supporting materials for this manuscript include the following:**

Datasets S1 to S5
Full gels and blots for Figures S4A, S5B, and S6A can be viewed at
https://figshare.com/s/89a474f54c67f67be215

**Supporting Information Text**

**SI Methods**

**Detailed cloning procedures**

**GFP-mCherry dual reporters.** An ~700 bp *mCherry* fragment was amplified by PCR from pDBP4164 using primers 62-53 and 62-54, and an ~800 bp *gfp* fragment was amplified from pDBP745 using primers 62-51 and 62-52 (oligonucleotides and strain genotypes are listed in **Table S1**). These fragments were fused by overlapping-PCR to generate a ~1.5 kb fragment which was ligated into Zero Blunt and stored as pDBP754 (*mCherry*WT). To generate *mCherry*$^{M71E}$ and *mCherry*$^{M71L}$, inverse PCR from pDBP754 was done using primer pairs 63-5 and 63-7 (*mCherry*$^{M71E}$) and 63-6 and 63-7 (*mCherry*$^{M71L}$) resulting in two ~5 kb fragments. These fragments were ligated into Zero Blunt and stored as pDBP760 (*mCherry*$^{M71E}$) and pDBP763 (*mCherry*$^{M71L}$). Plasmid constructs were verified by sequencing.

**p*tcu1::GFP-mCherry* dual reporters.** GFP-mCherry fragments (~1.8 kb) were amplified from plasmids pDBP754, pDBP760, and pDBP763 (this study) using primers 63-42 and 63-43. Fragments (~1 kb) upstream and downstream of the *tcu1* coding sequence were generated using WT (FGSC #4200) genomic DNA using primers 63-40, 63-41, 63-44, and 63-35. The three fragments were fused by overlapping-PCR to generate ~3.8 kb fusions that were ligated into Zero Blunt, verified by sequencing, and transformed into DH5α cells, and stored as plasmids pDBP794, pDBP796, pDBP797. The inserts were excised using EcoRI-HF and cotransformed with *csr1::hph*, which was excised out of pDBP429 using BssHII, into *Δmus-51::bar* (FGSC #9718) conidia and grown on plates containing 200 µg/mL Hygromycin B. Colonies were screened by GFP signal and native integration by PCR (63-82 and 63-83). Positive strains were backcrossed to *N. crassa* Δ*frq::bar* (DBP1228) and Δ*mak1::hph* (DBP835), and progeny expressing GFP-mCherry were stored as WT (DBP4480, DBP4641, and DBP4562), Δ*frq::bar* (DBP4484, DBP4496, and DBP4565), or Δ*mak1::hph* (DBP4487, DBP4505, and DBP4627).

**MetRS::LUC**. Fragments (~1 kb) upstream and downstream of the MetRS 3' end were generated from WT (FGSC #4200) using primers 54-28, 54-34, 54-37, and 54-33. A ~1.6 kb *luc* fragment was generated from pDBP366 using primers 54-35 and 54-36. The fragments were fused by overlapping PCR, ligated into Zero Blunt, the plasmids sequenced, and then transformed into DH5α cells, and stored as pDBP708. The MetRS::LUC insert was excised using EcoRI-HF and cotransformed with *csr1::hph*, which was excised out of pDBP429 using BssHII, into *Δmus-51::bar* (FGSC #9718) conidia and grown on plates containing 200 µg/mL Hygromycin B. Colonies were screened by luciferase signal and MetRS::LUC integration was verified using primers 54-38 and 54-39. Positive colonies were backcrossed to *N. crassa* Δ*frq::bar* (DBP1228) and progeny expressing luciferase were stored as MetRS::LUC (DBP4257) and MetRS::LUC, Δ*frq* (DBP4261).

**MetRS T108A and T108D mutations.** A silent mutation removing an XhoI restriction site was introduced into *metRS* using WT (DBP4278) *N. crassa* cells and primers 54-28, 65-5, 65-6, and 54-33. Next, *metRS* Thr 108 (ACT) to Ala 108 (GCA) (T108A) and Thr 108 (ACT) to Asp (GAT) (T108D) mutations were generated by overlapping-PCR using primers 54-28, 65-13 to 65-16, and 54-33. The ~4.8 kb *metRS* mutant fragments were ligated into Zero Blunt, sequenced, and then transformed into DH5α cells and stored as pDBP825 (*metRS*$^{T108A}$) and pDBP823 (*metRS*$^{T108D}$), respectively. T108A was excised from pDBP825 using EcoRI-HF and cotransformed with *csr1::hph*, which was excised out of pDBP429 using BssHII, into *Δmus-51::bar* (FGSC #9718) conidia and grown on plates containing 200 µg/mL Hygromycin B. Colonies were screened by PCR/digest of genomic DNA using primers 54-38 and 54-39 and XhoI and mutant strains were validated by genomic DNA sequencing. Positive transformants were backcrossed to WT (DBP4278), *ras-1*$^{bd}$ (DBP368), FRQ::LUC (DBP1563), *mCherry*WT (DBP4480), *mCherry*$^{M71E}$ (DBP4641), and *mCherry*$^{M71L}$ (DBP4562), and progeny were stored as WT (DBP4771), *ras-1*$^{bd}$ (DBP4932), FRQ::LUC (DBP4886), *GFP-mCherry*WT (DBP4802), *GFP-mCherry*$^{M71E}$ (DBP4799), and *GFP-mCherry*$^{M71L}$ (DBP4805).

*metRS*$^{T108D}$ was excised from pDBP823 using EcoRI-HF and cotransformed with pDBP429 into Δ*mus-51::bar* (DBP641), *GFP-mCherry*WT, Δ*mus-51::bar* (DBP5184), *GFP-mCherry*$^{M71E}$, Δ*mus-51::bar* (DBP5187), and *GFP-mCherry*$^{M71L}$, Δ*mus-51::bar* (DBP5188). The T108D mutation adds an Nru1 restriction site in the native *metRS* gene, and the mutation was screened by PCR/digest of genomic DNA using primers 54-72 and 66-54 and Nru1 and sequenced. *metRS*$^{T108D}$ positive colonies were stored as DBP4920, DBP5233, DBP5236 and DBP5239.

***bar::*p*tcu1::metRS::V5.*** A ~1.6 kb *V5::hph* was PCR-amplified from pDBP525 using primers 54-30 and 54-31. Homology arms (~1kb) were PCR-amplified from *bar::*p*tcu1::metRS* (DBP4059) using primers 54-28, 54-29, 54-32, and 54-33. Overlapping, split-marker PCR was used to separate the *hph* gene using primers 54-28 and 10-1 and 53-34 and 10-2. The two split-marker fragments were then transformed into *bar::*p*tcu1::metRS* (DBP4059) and cells were grown on plates containing 200 µg/mL Hygromycin B. Colonies were screened for native integration using primers 54-38 and 54-39. Colonies were backcrossed to WT (DBP4278) and progeny were screened by Hygromycin B resistance and stored as DBP4079.

**MetRS::CPD.** The *metRS* fragment (~2 kb ) was PCR-amplified from *N. crassa* cDNA using primers 63-29 and 63-30. The ~600 bp cysteine protease domain (CPD) gene was PCR-amplified from pDBP771 using primers 63-31 and 63-32, and both fragments were fused by overlapping PCR. NdeI-HF and NotI-HF restriction sites were added to the 5' and 3' ends using 63-46 and 63-47. MetRS::CPD and an *Escherichia coli* expression vector (pET-22b(+)) were digested using NdeI and Not1 and ligated to generate pDBP800, which was sequenced to validate.

**SI Figures**

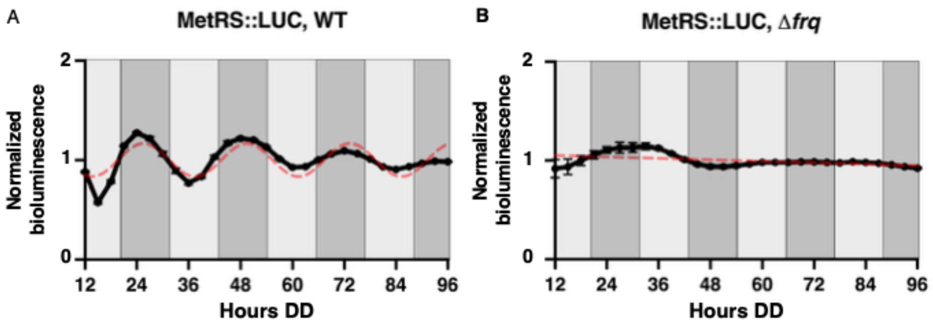


**Fig. S1. MetRS levels are clock-controlled.** Plot of bioluminescence levels from a MetRS::LUC translational fusion in WT (left) and clock mutant Δ*frq* (right) cells grown in DD over the indicated time (Hours DD). A) MetRS::LUC was rhythmic in WT cells as indicated by a better fit of the data to a sine wave (dotted red line; $p < 0.0001$) with a period of 23.8 ± 0.18 h. B) In contrast, MetRS::LUC rhythms were abolished in Δ*frq* cells as indicated by better fit to a straight line (dotted red line; $p > 0.05$). Error bars represent the mean ± SEM (n=12)

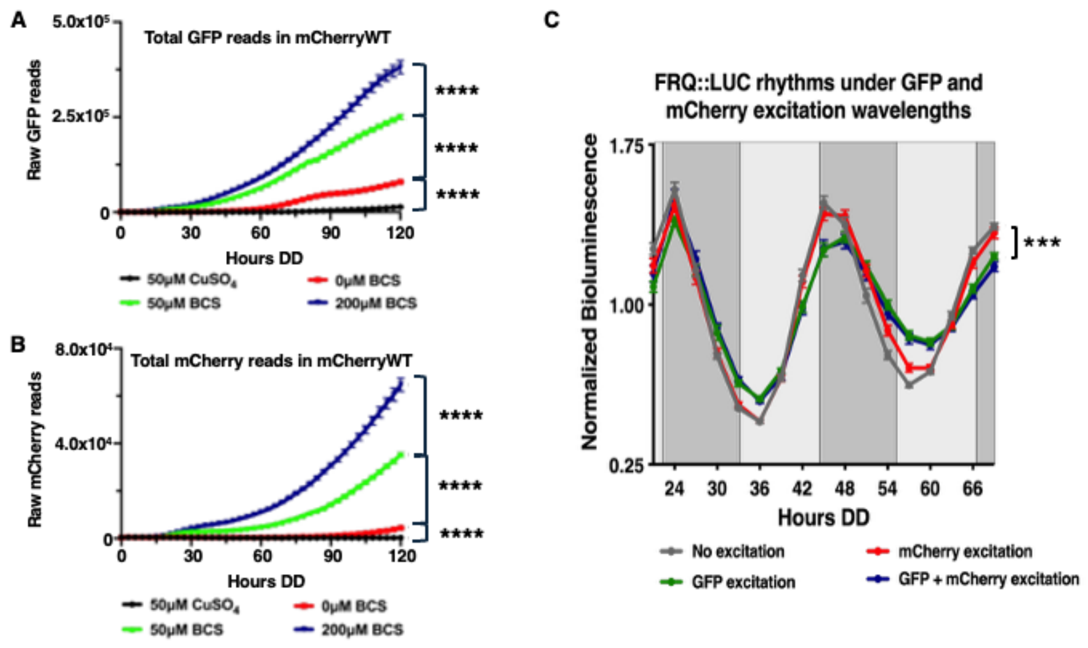


**Fig. S2. BCS drives constitutive expression of GFP-mCherry.** Plots of raw A) GFP and B) mCherry reads in GFP-mCherryWT. Cells were grown in DD at 25 °C with 50 µM $CuSO_4$ (black), 0 µM BCS (red), 50 µM BCS (green) or 200 µM BCS (blue) for 120 h. GFP and mCherry fluorescence were measured every three hours. Error bars represent the mean ± SEM (n = 6; ****$p < 0.0001$). C) Circadian rhythmicity persists upon GFP or mCherry excitation. Plot of FRQ::LUC rhythms under different excitation wavelengths. Cells were grown in DD at 25 °C and excited with mCherry (589nm) or GFP (488nm) wavelengths every three hours. Average bioluminescence is plotted, and error bars represent the mean ± SEM (n = 24). All plots are rhythmic; however, there was a significant decrease in amplitude upon GFP and GFP + mCherry excitation (***$p < 0.0005$).

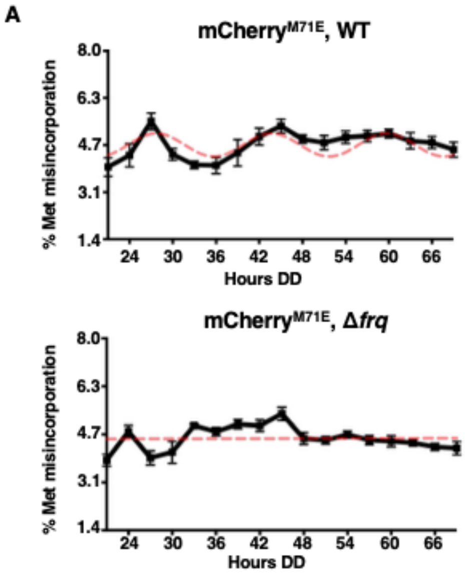


**Fig. S3. Met misincorporation in mCherry$^{M71E}$ is rhythmic in DD.** Plots of the percentage of Met misincorporation into A) mCherry$^{M71E}$ or B) mCherry$^{M71L}$ in WT and Δ*frq* cells grown in DD at 25 °C over 2 days (Hours DD). A) In WT cells (top), Met misincorporation in mCherry$^{M71E}$ peaked during the subjective night (DD28) and was rhythmic as indicated by a better fit of the data to a sine wave (dotted red line; $p < 0.005$). The rhythm was abolished in Δ*frq* cells (bottom) as indicated by better fit to a straight line (dotted red line, $p>0.05$). B) Met misincorporation was arrhythmic in mCherry$^{M71L}$ in both WT (top) and Δ*frq* (bottom) cells (dotted red line; $p > 0.05$). In all plots, error bars represent the mean ± SEM (n = 4).

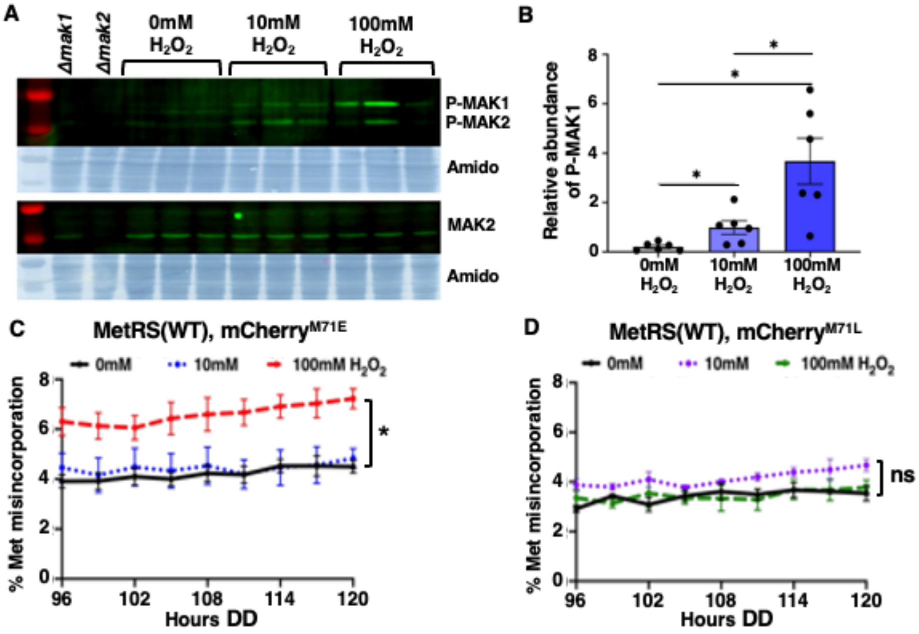


**Fig. S4. Oxidative stress induces P-MAK1 levels.** A) Representative western blot of three replicate samples (bracketed) from WT cells grown in constant light (LL) at 30 °C for 24 h and treated with 0 mM, 10 mM, or 100 mM $H_2O_2$ for 5 min and probed with anti-P-MAK1 and anti-MAK2 antibodies. MAK1 (Δ*mak1*) and MAK2 (Δ*mak2*) deletion strains served as controls for the antibodies. The anti-P-MAK1 antibody detects both P-MAK1 and P-MAK2, whereas the total anti-MAK2 antibody detects only MAK2. Amido black-stained gels served as protein loading controls. B) Plot of relative abundance of P-MAK1. P-MAK1 signals were normalized to total MAK2 to account for MAPK abundance/loading differences. Error bars represent the mean ± SEM (n = 6). Plots of percent Met misincorporation in cells grown in DD expressing C) mCherry$^{M71E}$ or D) mCherry$^{M71L}$ with or without treatment of different concentrations of $H_2O_2$ to induce oxidative stress. Measurements of Met misincorporation were taken at the indicated times (Hours DD) post-$H_2O_2$ treatment. Error bars represent the mean ± SEM (n = 3). In all plots, $*p < 0.05$, whereas ns represents no significant change.

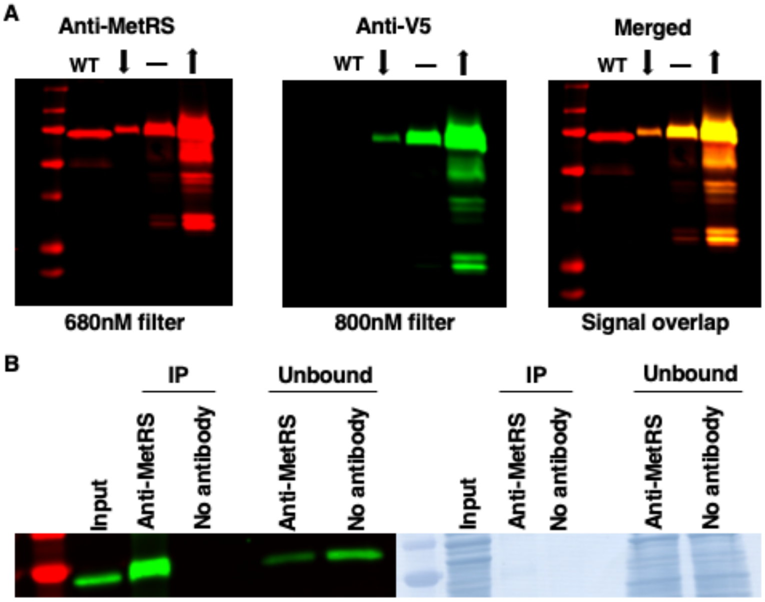


**Fig. S5. Anti-MetRS antibody is functional in western blots and immunoprecipitation.** A) Western blots of protein extracts from p*tcu1::metRS::V5* cells probed with rabbit anti-MetRS and and mouse anti-V5, and visualized under a 680 nm filter, 800 nm filter, and merged fluorescence signals. The endogenous *metRS* promoter was replaced with p*tcu1* to modulate expression of MetRS::V5 using copper. Tissue was grown for 24 hours in LL at 30 °C with 200 µM BCS (up-arrow), 0 µM BCS (flat line), or 200 µM $CuSO_4$ (down-arrow). Membranes were multiplexed with rabbit anti-MetRS and mouse anti-V5 primary antibodies and anti-rabbit (λ = 680 nM, red) and anti-mouse (λ = 800 nM, green) secondary antibodies. Anti-V5 and anti-MetRS signals colocalized as seen with the merged channel. B) Western blot of WT protein extracts immunoprecipitated using anti-MetRS antibodies. Tissue was grown in LL at 30 °C for 24 h and then harvested. Total protein extracts were stored as input and then 2 mg of protein extract were immunoprecipitated overnight at 4 °C with beads either bound to the anti-MetRS antibody (IP) or a no-antibody control (Unbound). The left panel is the western blot probed with anti-MetRS antibody, and the right panel is the corresponding amido black stain. MetRS was enriched in the IP sample using anti-MetRS antibody.

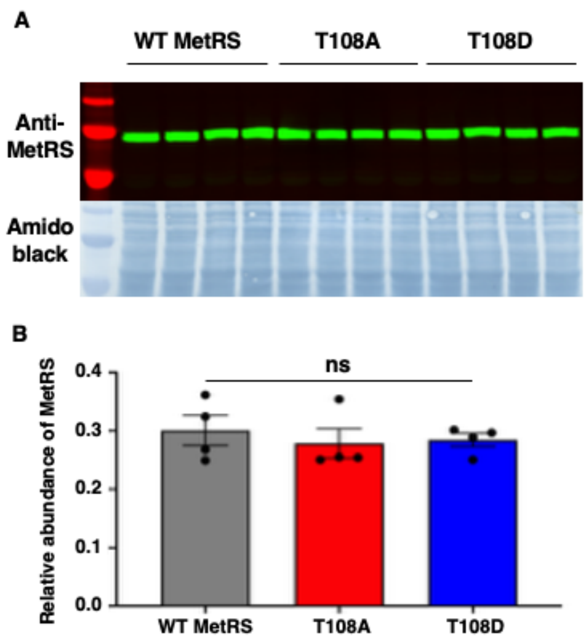


**Fig. S6. MetRS mutations do not affect total MetRS levels.** A) Western blot of 4 biological replicate protein extracts from WT, *metRS*$^{T108A}$ (T108A) and *metRS*$^{T108D}$ (T108D) cells grown for 24 hours in LL at 30 °C and probed with anti-MetRS antibody. Amido black-stained protein served as a loading control. B) Plot of normalized total MetRS levels of WT, *metRS*$^{T108A}$ (T108A) and *metRS*$^{T108D}$ (T108D) cells. There was no significant difference in MetRS abundance ($p > 0.05$). Error bars represent the mean ± SEM (n = 4; ns indicates no significant difference).

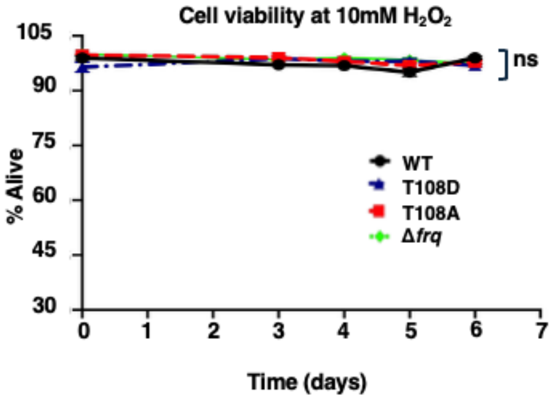


**Fig. S7. Cell viability is maintained in 10 mM $H_2O_2$.** Plot of cell viability for WT (black), *metRS*$^{T108A}$ (T108A, red), *metRS*$^{T108D}$ (T108D, blue) and Δ*frq* (green) cells grown in 10 mM $H_2O_2$. Cell viability was measured at zero-, three-, four-, five-, and six- days post-treatment. Error bars represent the mean ± SEM (n = 4; ns represents no significant change).

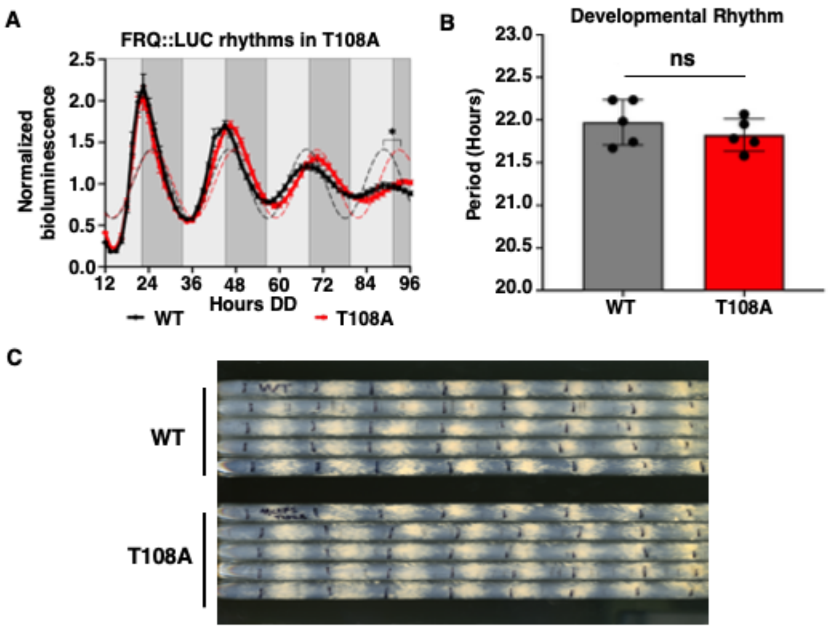


**Fig. S8. MetRS-phosphorylation is not essential for circadian clock function.** A) Plot of normalized FRQ::LUC expression in WT (gray) or *metRS*$^{T108A}$ (T108A, red) grown in DD 25 °C over the indicated time (Hours DD). Average bioluminescence signals were normalized and plotted. Error bars represent the mean ± SEM (n = 12). A one-hour lengthening of the period was observed in *metRS*$^{T108A}$ (T108A) compared to WT (*p < 0.05). B) Plot of period of the developmental rhythm in WT (gray) and *metRS*$^{T108A}$ (T108A, red). Error bars represent the mean ± SEM (n = 5). There was no significant difference (ns) in period between WT and *metRS*$^{T108A}$ (p > 0.05). C) Representative race tube assay demonstrating no change in the period of the circadian rhythm of conidiospore development in the *metRS*$^{T108A}$ (T108A) mutant compared to WT cells.

**SI Tables**

**Table S1: Met misincorporation in synonymous codons**

| Amino Acid Substitution | Codon | # Met Substitutions Observed | Genome Codon Count | Genome Fraction[1] | Expected[2] | Observed/ Expected |
|---|---|---|---|---|---|---|
| Glu to Met | GAG | 2071 | 253340 | 0.643 | 1696 | 1.22 |
| Glu to Met | GAA | 566 | 130563 | 0.357 | 940 | 0.50 |
| Asp to Met | GAC | 787 | 181331 | 0.570 | 728 | 1.08 |
| Asp to Met | GAT | 491 | 137024 | 0.430 | 550 | 0.89 |
| Leu to Met | CTC | 92 | 147846 | 0.320 | 94 | 0.98 |
| Leu to Met | CTG | 59 | 103055 | 0.220 | 66 | 0.90 |
| Leu to Met | CTT | 57 | 80868 | 0.170 | 52 | 1.11 |
| Leu to Met | TTG | 52 | 86881 | 0.185 | 55 | 0.94 |
| Leu to Met | CTA | 26 | 34153 | 0.070 | 22 | 1.20 |
| Leu to Met | TTA | 13 | 16791 | 0.036 | 11 | 1.21 |
| Total | | 4214 | | | 4214 | |

[1] Genome fraction is calculated as the codon count divided by the sum of all codons (365403 for Glu, 318356 for Asp, and 469594 for Leu).

[2] Expected number of Met substitutions is calculated by multiplying the genome fraction by the number of observed Met substitutions (2637 for Glu, 1278 for Asp, and 299 for Leu).

**Table S2.** Resources

| REAGENT or RESOURCE | RESOURCE | IDENTIFIE R |
|---|---|---|
| **Antibodies** | | |
| Rabbit polyclonal anti-MetRS (NCU07451) serum 1:10,000 | This paper | Anti-MetRS Serum 39087 |
| Rabbit monoclonal anti-P-p44/42 1:1,000 | Cell Signaling Technologies | Cat #9101S |
| Rabbit monoclonal anti-p44/42 1:1,000 | Cell Signaling Technologies | Cat #4695T |
| Mouse monoclonal anti-V5 1:5,000 | Invitrogen | Cat #R960-25 |
| Anti-rabbit 800nm LiCor IR dye 1:20,000 | Li-Cor | Cat #926-32219 |
| Anti-mouse 680nm LiCor IR dye 1:20,000 | Li-Cor | Cat #926-68070 |
| **Plasmids** | | |
| Plasmid PCR-blunt MetRS::LUC | This paper | pDBP708 |
| Plasmid ET MetRS::CPD | This paper | pDBP800 |
| Plasmid pRS416-10xGly::GFP | GenBank | Accession #AY598428 pDBP745 |
| Plasmid PCR-blunt *GFP-mCherry*(WT) | This paper | pDBP754 |
| Plasmid PCR blunt *GFP-mCherry*$^{M71E}$ | This paper | pDBP760 |
| Plasmid PCR blunt *GFP-mCherry*$^{M71L}$ | This paper | pDBP763 |
| Plasmid PCR-blunt p*tcu1::GFP-mCherry*(WT) | This paper | pDBP794 |
| Plasmid PCR-blunt p*tcu1::GFP- mCherry*$^{M71E}$ | This paper | pDBP796 |
| Plasmid PCR-blunt p*tcu1::GFP- mCherry*$^{M71L}$ | This paper | pDBP797 |
| Plasmid PCR-blunt MetRS$^{T108D}$ | This paper | pDBP823 |
| Plasmid PCR-blunt MetRS$^{T108A}$ | This paper | pDBP825 |
| Plasmid PCR-blunt *adv-1::V5::hph* | This paper | pDBP525 |

| REAGENT or RESOURCE | RESOURCE | IDENTIFIER |
|---|---|---|
| Plasmid Bluescript II KS (-) *csr1::hph* | (1) | pDBP429 |
| Plasmid BM61 Luciferase | (2) | pDBP366 |
| Plasmid PCR-blunt CPC-3::CPD antigen | This paper | pDBP771 |
| pET-22b(+) bacterial expression vector | Sigma-Aldrich | Cat #69744-3 |
| DH5α chemically competent *E. coli* | ThermoFisher | Cat #18265017 |
| Zero Blunt PCR cloning kit | Invitrogen | Ref #451245 |
| **Chemicals, peptides, and recombinant proteins** | | |
| Hygromycin B | VWR | Cat #80055-268 |
| Breathe-Easy membrane | Millipore | Cat #Z380059-1PAK |
| Bathocuproninedisulfonic acid (BCS) | Sigma-Aldrich | Cat #B1125 |
| Copper (II) sulfate ($CuSO_4$) | Sigma-Aldrich | Cat #C7631-250G |
| Hydrogen peroxide solution | Sigma-Aldrich | Cat #216763-100ML |
| Glucose | VWR | Cat #77985-508 |
| Revvity Optiplate-96 | Revvity | Cat #6005040 |
| 0.4% Trypan Blue | Thermo-Fisher | Cat #15250061 |
| Ni-NTA His Bind® Resin | Millipore | Cat# 70666-3 |
| Phytic acid solution ($InsP_6$) | Millipore | Cat# 83-86-8 |

| REAGENT or RESOURCE | RESOURCE | IDENTIFIE R |
|---|---|---|
| BssHII | New England Biolabs | Cat #R0199S |
| EcoRI-HF | New England Biolabs | Cat #R3101S |
| NotI-HF | New England Biolabs | Cat #R3189S |
| NdeI-HF | New England Biolabs | Cat #R0111S |
| NruI-HF | New England Biolabs | Cat #R3192S |
| **Strains** | | |
| *Neurospora crassa wild type* 74-OR23-IV mat a (WT) | FGSC | FGSC 4200, DBP4278 |
| Δ*mus-51::bar* | FGSC | FGSC 9718, DBP641 |
| Δ*frq::bar* | (3) | DBP1228 |
| Δ*mak1::hph* | (3) | DBP835 |
| Δ*mak2::hph* | (4) | DBP4343 |
| *ras-1*$^{bd}$ | (5) | DBP368 |
| *ccg-2::mCherry* | FGSC | FGSC10626 DBP4164 |
| p*tcu1::GFP::mCherry*(WT) | This paper | DBP4480 |
| p*tcu1::GFP::mCherry*$^{M71E}$ | This paper | DBP4641 |
| p*tcu1::GFP::mCherry*$^{M71L}$ | This paper | DBP4562 |
| p*tcu1::GFP::mCherry*(WT), Δ*frq::bar* | This paper | DBP4484 |
| p*tcu1::GFP::mCherry*$^{M71E}$, Δ*frq::bar* | This paper | DBP4496 |
| p*tcu1::GFP::mCherry*$^{M71L}$, Δ*frq::bar* | This paper | DBP4565 |
| p*tcu1::GFP::mCherry*(WT), Δ*mak1::hph* | This paper | DBP4487 |
| p*tcu1::GFP::mCherry*$^{M71E}$, Δ*mak1::hph* | This paper | DBP4505 |

| REAGENT or RESOURCE | RESOURCE | IDENTIFIE R |
|---|---|---|
| p*tcu1::GFP::mCherry*$^{M71L}$, Δ*mak1::hph* | This paper | DBP4627 |
| MetRS::LUC | This paper | DBP4257 |
| MetRS::LUC, Δ*frq::bar* | This paper | DBP4261 |
| FRQ::LUC | This paper | DBP1563 |
| *metRS::V5::hph* | This paper | DBP3953 |
| *bar::* p*tcu1::metRS* | This paper | DBP4059 |
| *bar::* p*tcu1::metRS::V5:hph* | This paper | DBP4079 |
| MetRS$^{T108A}$ | This paper | DBP4771 |
| *ptcu1::GFP::mCherry*(WT), Δ*mus-51::bar* | This paper | DBP5188 |
| *ptcu1::GFP::mCherry*$^{M71E}$, Δ*mus-51::bar* | This paper | DBP5187 |
| *ptcu1::GFP::mCherry*$^{M71L}$, Δ*mus-51::bar* | This paper | DBP5184 |
| MetRS$^{T108D}$ heterokaryon, Δ*csr1::hph,* Δ*mus-51::bar* | This paper | DBP4920 |
| p*tcu1::GFP::mCherry*(WT), MetRS$^{T108A}$ | This paper | DBP4802 |
| p*tcu1::GFP::mCherry*$^{M71E}$, MetRS$^{T108A}$ | This paper | DBP4799 |
| p*tcu1::GFP::mCherry*$^{M71L}$, MetRS$^{T108A}$ | This paper | DBP4805 |
| p*tcu1::GFP::mCherry*(WT), MetRS$^{T108D}$ heterokaryon, Δ*csr1::hph,* Δ*mus-51::bar* | This paper | DBP5233 |
| p*tcu1::GFP::mCherry*$^{M71E}$, MetRS$^{T108D}$ heterokaryon, Δ*csr1::hph,* Δ*mus-51::bar* | This paper | DBP5236 |
| p*tcu1::GFP::mCherry*$^{M71L}$, MetRS$^{T108D}$ heterokaryon, Δ*csr1::hph,* Δ*mus-51::bar* | This paper | DBP5239 |
| *ras-1*$^{bd}$, MetRS$^{T108A}$ | This paper | DBP4932 |
| FRQ::LUC, MetRS$^{T108A}$ | This paper | DBP4886 |
| **Oligonucleotides (purchased from IDT)** | | |
| To generate MetRS::LUC AF<br>5' – ACTCAAAGTCGCAACTGTCTC – 3' | This paper | OGB15<br>54-28 |
| To generate MetRS::LUC AR<br>5' – TGGCGTCCTCCCTCAAAGGAAGCTCCTTAG – 3' | This paper | OGB21<br>54-34 |
| To generate MetRS::LUC BF | This paper | OGB22 |

| REAGENT or RESOURCE | RESOURCE | IDENTIFIER |
|---|---|---|
| 5' -TCCTTTGAGGGAGGACGCCAAGAACATCAAG – 3' | | 54-35 |
| To generate MetRS::LUC BR<br>5' – CTAGTCTTTCTTAATCAGACGGCGATCTTG – 3' | This paper | OGB23<br>54-36 |
| To generate MetRS::LUC CF<br>5' – GTCTGATTAAGAAAGACTAGATATCCATTGGGTC – 3' | This paper | OGB24<br>54-37 |
| To generate MetRS::LUC CR<br>5' – TTGGTTGCCGAGTGAGAGAC – 3' | This paper | OGB20<br>54-33 |
| To verify MetRS construct integration AF<br>5' – CTGACGAAGAGTACGCCAAC – 3' | This paper | OGB25<br>54-38 |
| To verify MetRS construct integration AR<br>5' – TCCGTGTTCACTGCTCACAA – 3' | This paper | OGB26<br>54-39 |
| To generate MetRS::V5 AR<br>5' -CGATAAGCAACCTCAAAGGAAGCTCCTTAG – 3' | This paper | OGB16<br>54-29 |
| To generate MetRS::V5 BF<br>5' – TGGTTTGAGGTTGCTTATCGGCGGAGGCGG – 3' | This paper | OGB17<br>54-30 |
| To generate MetRS::V5 BR<br>5' – CTAGTCTTTCGTTTCACTTCCGAGCTCGGA – 3' | This paper | OGB18<br>54-31 |
| To generate MetRS::V5 CF<br>5' – GAAGTGAAACGAAAGACTAGATATCCATTGGGTC – 3' | This paper | OGB19<br>54-32 |
| To verify T108D integration<br>5' – AGAGATGCAGACGAACCAGG – 3' | This paper | OGB28<br>54-72 |
| To split *bar* cassette AR<br>5' – GTTGCGTGCCTTCCAGGGACC – 3' | (6) | P4 universal<br>13-78 |
| To split *bar* cassette BF<br>5' – GGAGACGTACACGGTCGACT – 3' | (6) | P5 universal<br>15-32 |
| To split *hph* cassette AR<br>5' – GTTGGTCAAGACCAATGCGGAGCA – 3' | (6) | Hy Rv<br>10-1 |

| REAGENT or RESOURCE | RESOURCE | IDENTIFIER |
|---|---|---|
| To split *hph* cassette BF<br>5' – CGACAGCGTCTCCGACCTGATG – 3' | (6) | Yg Fw<br>10-2 |
| To generate p*tcu1::GFP-mCherry* AF<br>5' – CGGAAACCGCCTGTCATTGCC – 3' | This paper | OGB95<br>63-40 |
| To generate GFP-mCherry AF<br>5' - ATGAGTATTCAACATTTCCGTGTCG - 3' | This paper | OGB75<br>62-51 |
| To generate GFP-mCherry AR<br>5' - GGTGGCGACTAGTGGATCCGACCAATGCTTAATCAGTGAGG - 3' | This paper | OGB76<br>62-52 |
| To generate GFP-mCherry BF<br>5' - TCGGATCCACTAGTCGCCACCATGGTGAGCAAGGGCGAGGAG - 3' | This paper | OGB77<br>62-53 |
| To generate GFP-mCherry BR<br>5' - TTACTTGTACAGCTCGTCCATGC - 3' | This paper | OGB78<br>62-54 |
| To generate *GFP-mCherry(M71E)* AF<br>5' - TGGAGCCGTACAGGAACTGAGGGACAGGA - 3' | This paper | OGB84<br>63-5 |
| To generate *GFP-mCherry(M71L)* AF<br>5' - TGGAGCCGTACTCGAACTGAGGGACAGGA - 3' | This paper | OGB85<br>63-6 |
| To generate *GFP-mCherry* mutants AR<br>5' - AGGCCTACGTAAAGCACCCCGCCGACATC - 3' | This paper | OGB86<br>63-7 |
| To generate p*tcu1::GFP-mCherry* AR<br>5' – GAATACTCATGGTTGGGGATGTGTGTGCGA – 3' | This paper | OGB96<br>63-41 |
| To generate p*tcu1::GFP-mCherry* BF<br>5' – ATCCCCAACCATGAGTATTCAACATTTCCG – 3' | This paper | OGB97<br>63-42 |
| To generate p*tcu1::GFP-mCherry* BR<br>5' – GAGTGGAAGGTTACTTGTACAGCTCGTCCA – 3' | This paper | OGB98<br>63-43 |
| To generate p*tcu1::GFP-mCherry* CF | This paper | OGB99 |

| REAGENT or RESOURCE | RESOURCE | IDENTIFIE R |
|---|---|---|
| 5' – GTACAAGTAACCTTCCACTCAAACCCTGTA – 3' | | 63-44 |
| To generate p*tcu1::GFP-mCherry* CR<br>5' – CAGTGAATTCTACAAGAAGGACGATC – 3' | This paper | OGB100<br>63-45 |
| To verify integration of p*tcu1::GFP-mCherry*<br>5' – AGTCATAACAACCGGACCTC – 3' | This paper | OGB118<br>63-82 |
| To verify integration of p*tcu1::GFP-mCherry*<br>5' – AAGCTCAAGGATACATTCGG – 3' | This paper | OGB119<br>63-83 |
| To PCR MetRS out of cDNA for MetRS::CPD/antibody generation AF<br>5' – ACATATGGCGGCCGCATGGCCAGCAAAAACGGACC – 3' | This paper | OGB87<br>63-29 |
| To PCR MetRS out of cDNA for MetRS::CPD/antibody generation AR<br>5' – TTCCATCCGCCCTCAAAGGAAGCTCCTTAG – 3' | This paper | OGB88<br>63-30 |
| To PCR CPD out of cDNA for MetRS::CPD/antibody generation AF<br>5' – TCCTTTGAGGGCGGATGGAAAAATACTCCA – 3' | This paper | OGB89<br>63-31 |
| To PCR CPD out of cDNA for MetRS::CPD/antibody generation AR<br>5' – GTGGTGCTCGAGACCTTGCGCGTCCCAGCTTA – 3' | This paper | OGB90<br>63-32 |
| To remove XhoI site in MetRS for mutation screening AR<br>5' – ACCCCCGAGGCCCTTGGGGAC – 3' | This paper | OGB140<br>65-5 |
| To remove XhoI site in MetRS for mutation screening BF<br>5' – GTCCCCAAGGGCCTCGGGGGT – 3' | This paper | OGB141<br>65-6 |
| To add 5'-NdeI and 3'-NotI sites to MetRS::CPD AF<br>5' – GTGGTGCATATGGCCAGCAAAAACGGACC – 3' | This paper | OGB101<br>63-46 |
| To add 5'-NdeI and 3'-NotI sites to MetRS::CPD AR<br>5' – GCGGCCGCACCTTGCGGCGTCCCAGCTTA – 3' | This paper | OGB102<br>63-47 |
| To generate T108A mutation AR<br>5' – TTCGGTCGCGATCCCACCGA – 3' | This paper | OGB142<br>65-13 |

| REAGENT or RESOURCE | RESOURCE | IDENTIFIER |
|---|---|---|
| To generate T108A mutation BF<br>5’ – TCGGTGGGATCGCGACCGAA – 3’ | This paper | OGB143<br>65-14 |
| To generate T108D mutation AR<br>5’ – TTCGGTCGCGCACCCACCGA – 3’ | This paper | OGB144<br>65-15 |
| To generate T108D mutation BF<br>5’ – TCGGTGGGTGCGCGACCGAA – 3’ | This paper | OGB145<br>65-16 |
| To verify integration of MetRS$^{T108D}$ by NruI digestion with OGB26 AF<br>5’ – TGGGACGATGGATCCACTGC – 3’ | This paper | OGB184<br>66-54 |
| **Software and Algorithms** | | |
| Cosine wave analysis | (5) | GraphPad Prism software package |
| ECHO | (7) | |
| **Other** | | |
| Odyssey-CLX infrared imaging system | Li-Cor | Cat#CLX-2080 |
| EnVision Xcite Multilabel Reader with Monochromator add-on | PerkinElmer | Cat# 2105-0010 |

**SI Datasets**

**Dataset S1.**Raw data of MetRS immunoprecipitation/mass spectrometry with trypsin digestion to identify posttranslational modifications from WT, ∆*mak1* and ∆*mak2* cells with or without $H_2O_2$ treatment.

**Dataset S2.** Raw data of Andromeda (8) search for Met misincorporated peptides in WT or Δ*csp1* from a published rhythmic proteomics dataset (9), related to Figure 5

**Dataset S3.** Gene ID lists for proteins with identified Met misincorporated peptides in WT or Δ*csp1*, related to Figure 5 and Dataset S2.

**Dataset S4.** Raw data of MS-GF+ search for Met misincorporated peptides in WT cells from a published rhythmic proteomics dataset, related to Figure 5.

**Dataset S5.** ECHO (7) outputs and GO analysis for all rhythmic Glu to Met misincorporated peptides in WT or Δ*csp1*, related to Figure 5 and Dataset S4.
